# Experimental Framework to Investigate Glioma in Organotypic Human Cortex

**DOI:** 10.64898/2026.09.21.753089

**Authors:** Lotte R. Woltheus, Lucas Baudouin, Marike van Lingen, Niels Verburg, Sander Idema, Myron G. Best, Phillip de Witt Hamer, David Noske, Linda Douw, Dorien Maas

## Abstract

**Background:** Gliomas are primary brain tumors that integrate into the surrounding brain via neuron-glioma synapses and disrupt normal neuronal functioning. Recent work links tumor-brain connectivity to clinical measures such as patients’ functional status. However, few experimental systems allow the visualization of tumor and resident brain cell architecture together in human tissue at the scale needed to connect cellular organization to clinical outcomes. To address this gap, we developed an experimental framework for investigating the neuron-glioma network in organotypic human cortex from glioma patients, designed to support future correlation of cellular architecture to clinical outcomes such as functional status and survival.

**Methods:** Brain tissue samples containing cortex from seven glioma patients were collected during tumor resection and cultured for seven days as organotypic slices. We established several multiplexed immunohistochemistry panels to stain for glioma cells, various types of resident brain cells including neurons and cells from the oligodendrocyte lineage, as well as axonal networks and myelination patterns.

**Results:** We demonstrated the presence of tumor cells in all samples using SOX2, a tumor cell marker. NeuN+ neuronal cells were also successfully identified, as were cells from the oligodendrocyte lineage using SOX10 (pan-lineage marker), BCAS1 (pre-myelinating oligodendrocyte marker), and ASPA (mature oligodendrocyte marker). The neuronal network and myelin were successfully visualized using MAP2 (neurons and dendrites), SMI312 (axons), and PLP1 (myelin).

**Conclusions:** High-quality visualization of tumor cells, resident brain cells and the (myelinated) neuronal network in organotypic human cortex from glioma patients is possible after seven days of culturing. Future studies could further optimize culture conditions and should study stability of all cellular components in longer term slice cultures. Preliminary analyses indicate that cellular measures derived from human organotypic slices can be linked to patients’ functional status, demonstrating the capacity to generate data suitable for large-scale investigation of the effects of glioma-brain network architecture on clinical outcomes.

## Introduction

Gliomas are primary brain tumors that cause a variety of symptoms, such as seizures, cognitive impairment, and motor deficits, at the time of diagnosis and beyond [1]. Symptoms can affect patients’ functional status and independence, which is assessed by clinicians using the Karnofsky Performance Status (KPS) scale. KPS is used not only in clinical settings but also extensively in research as a clinical outcome. However, KPS is only partly explained by conventional tumor features, such as location and size [2]. Understanding how glioma influences brain function and, subsequently, functional status remains a challenge.

Based on recent research, the functional status of glioma patients is associated with tumor-brain integration [2-4]. Gliomas are not isolated tumors but are embedded into the brain tissue and diffusely interacting with their surrounding environment [5,6]. Glioblastoma (GBM) is the most aggressive form of glioma, and research on integration of gliomas into the brain tissue is mostly done in this subtype. GBM cells are known to be interconnected through microtubes. Microtubes facilitate tumor invasion by connecting tumor cells via gap junctions to form a functional tumor network in which autonomous cell activity is easily propagated throughout the tumor [5,6]. However, this microtube-based network varies across glioma subtypes [5]. In addition to intratumoral connections, GBM cells also form synapses with neurons. Different subtypes of neuron-glioma synapses have been identified, including glutamatergic [7], GABAergic [8], cholinergic [9], and serotonergic [10] synapses. Synaptic activation induces rhythmic cell activity within the tumor cell network [7]. The neuron-glioma synapses create a positive feedback loop where glioma cells enhance neuronal excitability, and neuronal activity, in turn, promotes glioma proliferation and secretion of factors that increase excitability [11,12]. GBM cells thus form a complex network that is functionally integrated into neuronal networks, and that influences brain activity and may be a critical determinant of patients’ functional status. It remains to be seen whether other forms of glioma such as IDH-mutant gliomas form the same kinds of synaptic contacts and integration into the brain network.

Next to functional integration of tumor cells into the brain network, structural integration into the brain’s white matter has also been reported. Glioma tends to occur in highly structurally embedded brain regions [3]. In mouse models, tumor-infiltrating cells have been shown to infiltrate the cortex along white matter tracts [13,14]. White matter tracts consist of myelinated axons, and myelin is an extension of the oligodendrocyte cell membrane. Oligodendrocyte lineage cells have been found to communicate bidirectionally with glioma cells [15]. Also, in a mouse model, GBM infiltration disrupts myelination [16]. Similarly, disrupted myelin morphology associated with GBM, was observed in a study combining diffusion MRI and immunohistochemistry. This altered white matter morphology correlated with poorer pre-operative muscle strength compared to other white matter structures typical of lower-grade glioma [17]. Thus, glioma cells integrate into and disrupt white matter structures. However, how the organization of this structural architecture influences the tumor-brain network and its association with the KPS score and whether this differs in glioma subtypes remains unknown.

Developing an experimental paradigm that allows investigation of the role of the structural and functional embedding of gliomas into the brain is important to investigate relationships with functional outcomes such as KPS and glioma subtype differences. Studying this relationship requires an experimental model that simultaneously preserves tumor architecture and interactions with resident brain cells, while allowing functional measurements. Two-dimensional cellular cultures lack the tumor microenvironment that is crucial for examining the tumor architecture. An *in vivo* mouse model would provide the architecture but would not comprehensively mimic human-specific glioma behavior [18]. Therefore, a human organotypic slice culture model would be ideal for such investigations. Using organotypic human cortex from glioma patients offers several advantages, including preservation of the tumor microenvironment and cytoarchitecture [19,20]. Additionally, observations across scales can be better understood and put into a patient-specific context. As such, organotypic slice cultures are increasingly used in glioma research. However, studies vary significantly in their applied approach [19]. Altogether, organotypic human cortex slice cultures appear to be the ideal model, but standardization of protocols is needed.

In this study, the aim was therefore to establish an experimental framework for organotypic slice cultures from resected human glioma tissue with replicable outcome measures that allow visualization and quantification of tumor cells and resident brain cells in subtypes of glioma. Immunohistochemical staining panels visualizing tumor cells, neurons, axons, dendrites, myelin, and oligodendrocyte subpopulations were developed. Moreover, preliminary analyses were conducted, showing the feasibility of connecting cellular features to KPS. Ultimately, by creating this human organotypic slice culture analysis framework, we provide valuable research tools that may help to deepen our understanding of the relationship between a patient’s functional status and the underlying cellular architecture.

## Methods

### Clinical data and patient-derived organotypic slice cultures

This study involved collecting brain tissue from glioma patients at the Amsterdam University Medical Center (Amsterdam UMC). To gain access to a subcortically located tumor, the surgeon resected cortical brain tissue that was ‘en route’ towards the tumor and was therefore considered waste material. The resected cortical samples were obtained with sharp dissection and without tissue coagulation to reduce artifacts and were subsequently used for culturing and experiments. All patients gave written informed consent for the use of their medical data as obtained through standard clinical care and leftover tissue prior to participation (Medical Ethics Committee of VU University Medical Center reference 2010/126) according to the Declaration of Helsinki. Patient characteristics can be found in Table 1. IDH-status was determined post-operatively through histopathological examination and molecular testing of the resected brain tumor in accordance with the World Health Organization 2021 classification [21]. Additionally, the pre-operative Karnofsky Performance Status (KPS) was recorded for each patient by their treating physician. To compare high and low KPS scores, patients were subdivided into two groups: KPS ≤ 80 and KPS > 80 [22].

**Table 1.** Patient characteristics.

| | KPS $\leq 80$ (n=3) | KPS $> 80$ (n=4) |
| --- | --- | --- |
| **Demographics** |  |  |
| Age at surgery (median, range) | 72 (36-73) | 38 (28-59) |
| Sex (male/female) | 1/2 | 2/2 |
| **Tumor characteristics** |  |  |
| IDH (wild-type/mutant) | 3/0 | 1/3 |
| Tumor grade (median, range) | 4 (4-4) | 3 (2-4) |
| Tumor location | Frontotemporal 1, Temporal 1, Parietooccipital 1 | Frontal 1, Frontal insular 2, Mesiotemporal 1 |
| Tumor laterality (left/right) | 2/1 | 3/1 |
| **Clinical information** |  |  |
| Pre-op KPS (range) | 70-70 | 90-100 |
| Epilepsy (yes/no) | 1/2 | 4/0 |
| AED use (yes/no) | 1/2 | 4/0 |

Upon surgical resection, the cortex samples were immediately kept in ice-cold N-Methyl-D-glucamine (NMDG) solution [23]. Slices of 250 or 350 µm thickness were cut using a vibratome (Leica, VT1200), in ice-cold carbonated NMDG solution. The sections were transferred into a Millicell® (Sigma Aldrich, PICM03050) within a 6-well plate containing pH- and temperature-equilibrated culturing medium. The medium was refreshed every 48 to 72 hours and consisted of Minimum Essential Medium (MEM; Gibco™, Thermo Fisher Scientific, 11575-032; 78.3% v/v) supplemented with ascorbic acid (0.84 mM; Sigma, A7631), HEPES (24.7 mM; Merck, H3537), D-glucose (10.3 mM; Gibco™, A2494001), penicillin/streptomycin (20.6 U/mL; Invitrogen, 15070063), GlutaMAX™ (0.21×; Gibco™, 35050061), MgSO_4_ (1.65 mM; Merck, M1880), CaCl_2_ (0.82 mM; Merck, 21115), bovine pancreas insulin (0.72 µM; Merck, I0516), and heat-inactivated horse serum (16.5%; Thermo Fisher Scientific, 26050088) [24]. The sections were incubated at 37 °C with 5% CO_2_ for 7 days.

### Sample preparation and immunohistochemistry

Sections were fixed for 1 hour with 2% paraformaldehyde and washed three times for 5 minutes with 1X phosphate buffer (PBS), either immediately after collection or after culturing for 7 days in vitro. For tissue preservation, samples were embedded in 10% sucrose (Sigma-Aldrich, 16104) in 1X PBS for at least 2 hours at 4 °C, then overnight in 20% sucrose, followed by another overnight in 30% sucrose. Sucrose-embedded samples were frozen and emerged in Optimal Cutting Temperature (OCT) compound (VWR®, 361603E). Samples were then resectioned in a CryoStar NX50 cryostat (Thermo Scientific) at 12, 14, or 20 µm and stored at -20 °C.

Before staining, slides were brought to room temperature and submerged in 1X TBS for 30 minutes. Next, slides were washed twice for 2 minutes with 1X TBS. Permeabilization and blocking of non-specific antibody binding were performed for 30 minutes in a 1X TBS solution containing 5% Donkey serum (Jackson, 017-000-121) and 0.4% Triton-100 (Merck, 8603). Slides were then incubated overnight at 4 °C with primary antibodies (Table 2) diluted in 1X TBS with 5% Donkey serum and 0.05% Triton-100. The following day, slides were washed three times for 5 minutes each with 1X TBS containing 0.05% Tween-20 (Sigma, P7949), then incubated with secondary antibodies (Table 2) and DAPI (1:1000; Sigma, D9542) for 30 to 60 minutes in 1X TBS with 5% Donkey serum and 0.05% Triton-100. After incubation, slides were washed three times for 5 minutes with 1X TBS containing 0.05% Tween-20, followed by a quick rinse in 1X TBS. Slides were then treated for 1 minute with TrueBlack Lipofuscin Autofluorescence Quencher (Biotium, 23007), diluted 1:50 in 70% ethanol. Finally, after two 5-minute washes with 1X TBS, slides were mounted with coverslips using Fluoromount G (Southern Biotech, 0100-01) and stored at 4 °C.

**Table 2.** Primary and secondary antibodies.

| Type | Antigen<br>/Fluorophore | Host<br>Species | Target<br>Species | Antibody | Identifier<br>(RRID) | Concentration |
| --- | --- | --- | --- | --- | --- | --- |
| Primary | SOX2 | Rabbit | - | AB5603<br>(Millipore) | AB_2286686 | 1:500 |
| Primary | NeuN | Mouse | - | MAB377<br>(Chemicon) | AB_2298772 | 1:500 |
| Primary | MAP2 | Chicken | - | NB300-213<br>(Novus) | AB_2138178 | 1:500 |
| Primary | PLP1 | Rabbit | - | 33901-1-AP<br>(ProteinTech) | AB_3743106 | 1:400 |
| Primary | SMI312 | Mouse | - | 837904<br>(BioLegend) | AB_2566782 | 1:500 |
| Primary | SOX10 | Goat | - | AF2864 (R&D) | AB_442208 | 1:500 |
| Primary | BCAS1 | Mouse | - | Sc-136342<br>(Santa Cruz) | AB_10839529 | 1:750 |
| Primary | ASPA | Rabbit | - | 13244-1-AP<br>(ProteinTech) | AB_2274358 | 1:500 |
| Primary | Nestin | Rabbit | - | AB5922<br>(Chemicon) | AB_91107 | 1:1000 |
| Primary | VGlut1 | Guineapig | - | 135304<br>(Synaptic<br>Systems) | AB_887878 | 1:500 |
| Primary | GriA1 | Mouse | - | 67642-1-Ig<br>(ProteinTech) | AB_2882842 | 1:1000 |
| Secondary | Alexa 488 | Donkey | Rabbit | A21206 (Mol.<br>Probes) | AB_2535792 | 1:500 |
| Secondary | Alexa 594 | Donkey | Mouse | A21203 (Mol.<br>Probes) | AB_141633 | 1:500 |
| Secondary | Alexa 488 | Donkey | Chicken | 703-545-155<br>(Jackson) | AB_2340375 | 1:1000 |
| Secondary | Alexa 546 | Donkey | Rabbit | A10040<br>(Invitrogen) | AB_2534016 | 1:1000 |
| Secondary | Alexa 647 | Donkey | Mouse | A32787<br>(Invitrogen) | AB_2762830 | 1:500 |
| Secondary | Alexa 488 | Donkey | Goat | A11055 (Mol.<br>Probes) | AB_2534102 | 1:1000 |
| Secondary | Alexa 546 | Donkey | Mouse | A10036 (Mol.<br>Probes) | AB_11180613 | 1:1000 |
| Secondary | Alexa 647 | Donkey | Rabbit | A32795<br>(Invitrogen) | AB_2762835 | 1:500 |
| Secondary | Alexa 488 | Donkey | Guineapig | 706-545-148<br>(Jackson) | AB_2340472 | 1:1000 |

### Image and data analysis

Imaging was performed using a slide scanner (SLIDEVIEW^TM^ VS200, Olympus/Evident) equipped with a 20x air objective (NA = 0.8; final resolution, 0.345 µm/pixel) and filters for DAPI, FITC, Cy3, and Cy5. Images were also taken using a confocal microscope (Leica TCS SP8 STED 3X, Leica Microsystems) with a 40x oil immersion objective (NA = 1.3; final resolution, 0.284 µm/pixel) for analysis of neuronal and myelin architecture. For each cryosection, the maximum intensity projection was generated. All images were processed using QuPath (v0.5.1), with sections manually delineated and further defined via pixel-based detection by averaging channel intensities to exclude tissue holes and artefacts. Cells were identified based on DAPI staining, with an intensity threshold of 120. SOX2 and NeuN positive cells were annotated using single-measure classifiers, with thresholds based on nuclear intensity standard deviation. Images were processed in ImageJ (v1.54f) for visualization and scale bar addition. Graphs were made using GraphPad Prism 10.6.0. Due to the small sample size, variations in cryosection thicknesses, and demographic differences between KPS groups, no statistical tests were conducted.

## Results

### Evaluation of organotypic slice culture preparation

A total of 22 organotypic slices from seven patients were collected at thicknesses of 250 µm or 350 µm (Fig 1a). Initially, from two patient samples, sections treated and untreated with a sucrose solution were compared for tissue preservation. This sucrose protection was deemed necessary, and subsequently, all samples were treated. Samples that were not treated were used for pilot stains and optimization. Additionally, the thicknesses of 250 µm and 350 µm were compared, and the 350 µm sections appeared more suitable for tissue integrity. In cases where the available tissue was relatively small, sections of 250 µm thickness were still collected. Ultimately, tissue samples from two out of seven patients did not progress successfully through the pipeline (Table 1). This was either due to initial sectioning issues or poor sample quality, possibly caused by mechanical stress during collection. Moreover, of the remaining five patient samples, some sections were successfully re-sectioned, while others were not successfully re-sectioned.

**Figure 1.**
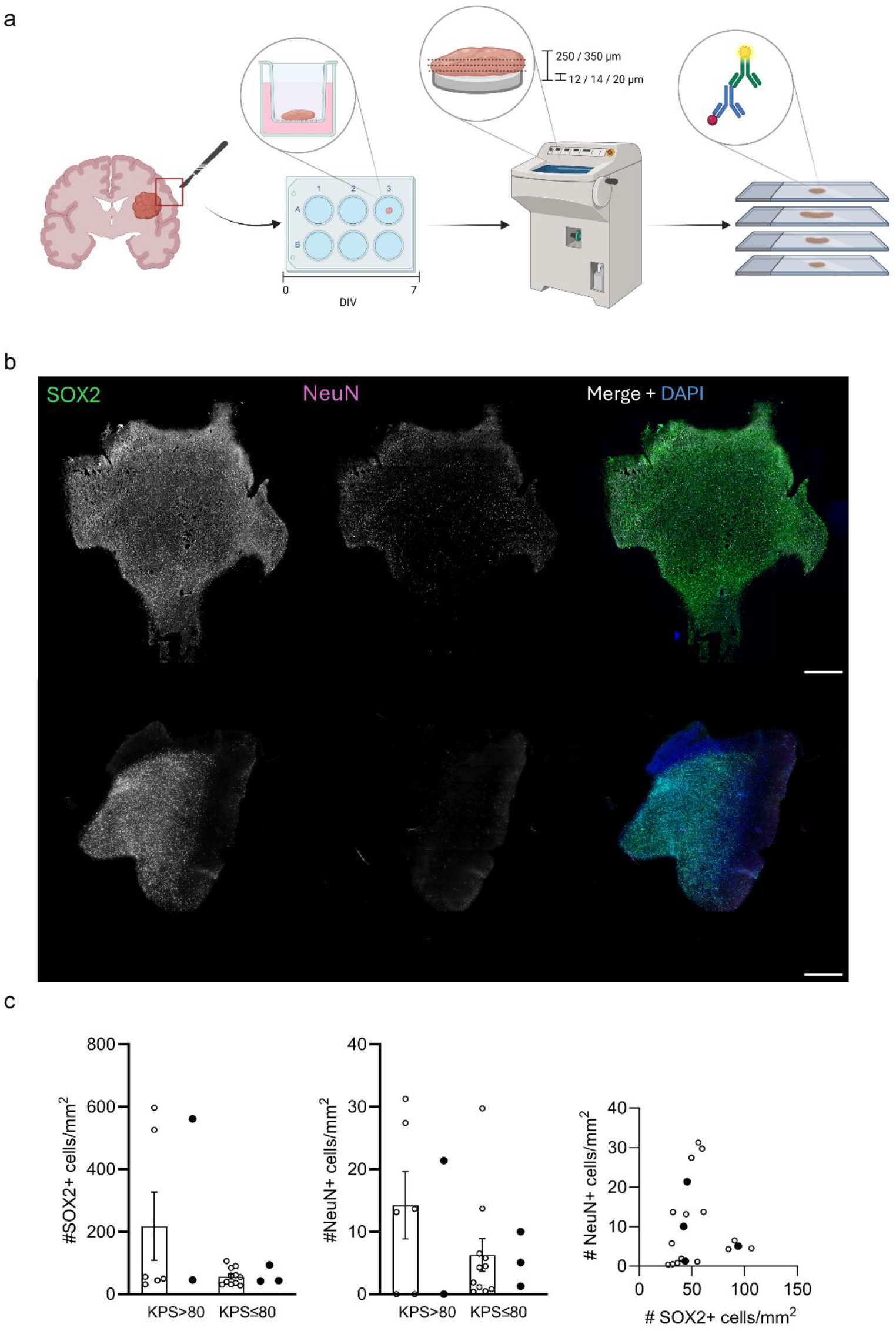
Visualization of SOX2+ glioma cells and NeuN+neurons in organotypic slices from glioma patients after 7 days *in vitro*. A) Experimental protocol included using resected cortex samples from glioma patients, culturing of organotypic cortex slices for seven days, cryo-sectioning and subsequent immunohistochemistry. DIV: day in vitro. Created with BioRender (https://core.local.biorender.dev/api/short-link/1jhuoeq). B) 20x maximum intensity projection images from fixed-frozen cryosections from a surgical cortex sample of an adult human glioma patient. The upper row includes an IDH wild-type sample (12 µm), and the bottom row includes an IDH mutant sample (20 µm). Staining for SOX2 and NeuN with a DAPI nuclear counterstain. Scale bars are 1 mm. C) Numbers of SOX2+ (left) or NeuN+ (middle) cells /mm2 as well as a correlation between these measures (left). Open dots represent values from cryosections, closed dots represent patient averages. The bargraphs represent average +/-standard error of the mean. In the correlation graph we included only values from sections cut at 12 µm. KPS: Karnofsky Performance Status.

### SOX2+ cell infiltration and NeuN+ cell quantity

Immunohistochemistry with a SOX2 antibody allowed the visualization of glioma cells in the organotypic cortex samples (Fig 1b). Glioma cell infiltration was calculated as the number of SOX2+ cells / mm^2^ per cryosection and averaged per patient [15]. All stained samples contained SOX2+ cells, although the number of positive cells varied (Fig 1c). Additionally, to assess the quantity of neurons, the samples were stained with NeuN (Fig 1b). Similarly to SOX2, the number of NeuN+ cells varied between samples (Fig 1c). Although the sample size did not allow for statistical comparisons, preliminary analyses indicate that cell density data can be linked to KPS scores and that correlations between cell densities can be calculated (Fig 1c).

Notably, all samples had low numbers of NeuN+ cells (14.25 +/-13.20 (mean +/-standard deviation) NeuN+ cells / mm^2^ in 11 organotypic slices from 3 patients with KPS > 80; 6.29 +/-8.70 NeuN+ cells / mm^2^ in 6 organotypic slices from 2 patients with KPS ≤ 80). This low count of NeuN+ cells might be partly due to the effects of culturing [20]. To investigate these cultivation-dependent effects on the quantity of both SOX2+ and NeuN+ cells, an acute (uncultured) section was compared with a cultured section from a single patient. This comparison revealed a lower number of NeuN+ and SOX2+ cells in the cultured sample compared to the acute section (Fig 2a). This discrepancy implies a cultivation-dependent impact on NeuN expression or the number of NeuN-expressing cells. While differences in SOX2+ cell numbers might also reflect variations in infiltration. Overall, these observations indicate that glioma infiltration can be visualized in organotypic cortex samples, although the number of NeuN+ cells appears reduced after seven days of culture.

**Figure 2.**
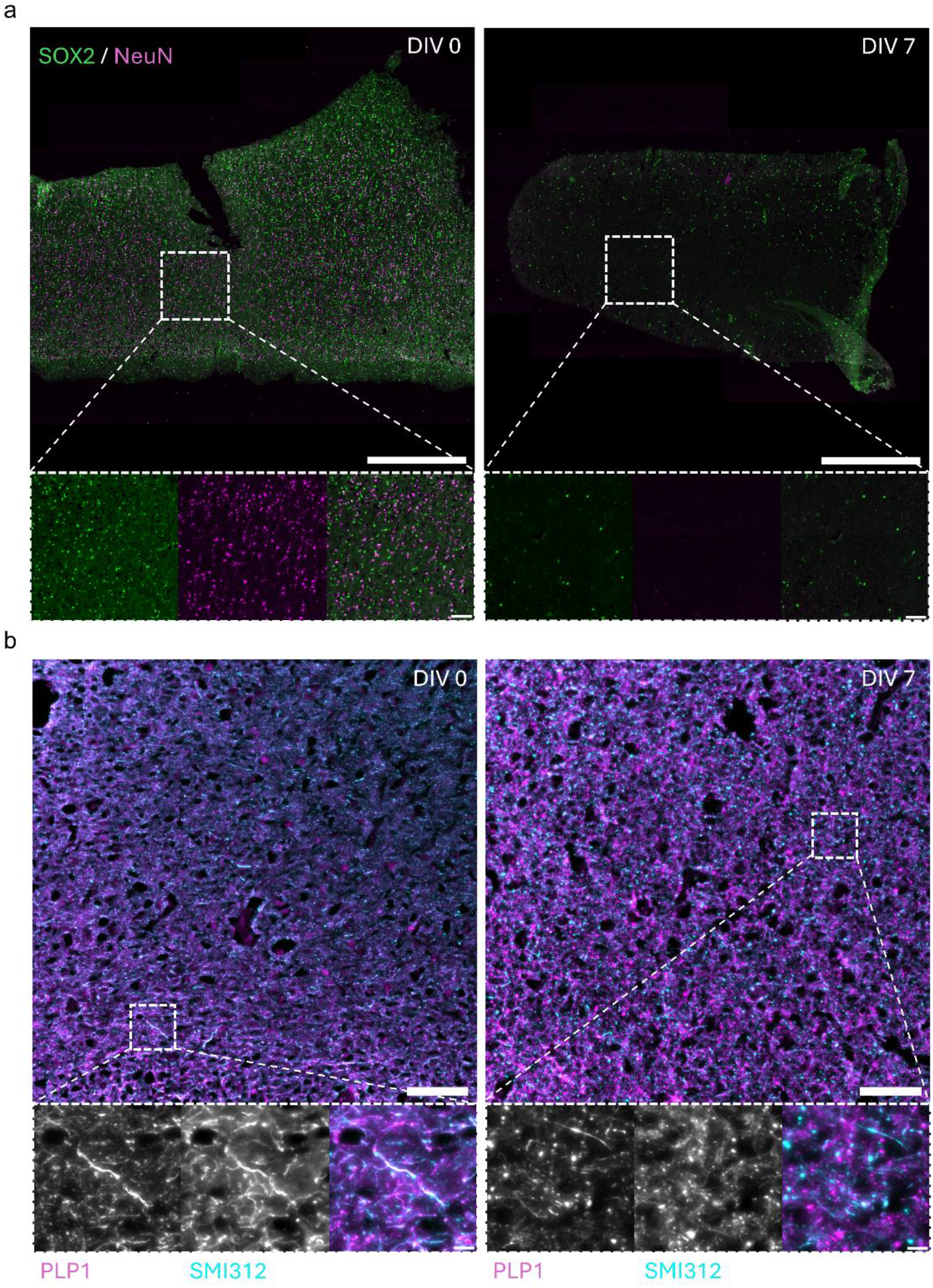
Culture-dependent effects on glioma and neuron numbers and myelin organization. Acute and DIV7 fixed-frozen cryosections (12 µm) originating from the same surgical cortex sample from an adult human glioma patient; IDH wild-type tumor. Maximum intensity projection images from a 20x Slide scanner. DIV: day in vitro. A) Staining including SOX2 and NeuN. Scalebars: 1 mm and 100 µm. B) Staining, including PLP1, and SMI312. Scalebars: 100 µm and 10 µm.

### Neuronal network composition and myelin visualization

For a more detailed examination of neuronal architecture and myelination, subsequent stains with MAP2, SMI312, and PLP1 antibodies were performed. Several DAPI-stained cells tested positive for MAP2, confirming their neuronal identity (Fig 3a). Additionally, axons were visualized through SMI312 and myelin through PLP1 (Fig 3a-b). Initially, an antibody targeting the heavy chain of neurofilament, NF-200 [25], was used; however, its application was unsuccessful after multiple trials. Conversely, SMI312, an antibody against the highly phosphorylated medium and heavy neurofilament subunits [26], proved successful. Variations in staining patterns and marker overlap were observed within sections. Overlapping glioma infiltration hotspots (based on SOX2+ cells from Fig 1b, upper row) with PLP1 coverage revealed specific dark spots coinciding with glioma cell infiltration (Fig 3b). These dark areas suggest disintegrated myelin.

**Figure 3.**
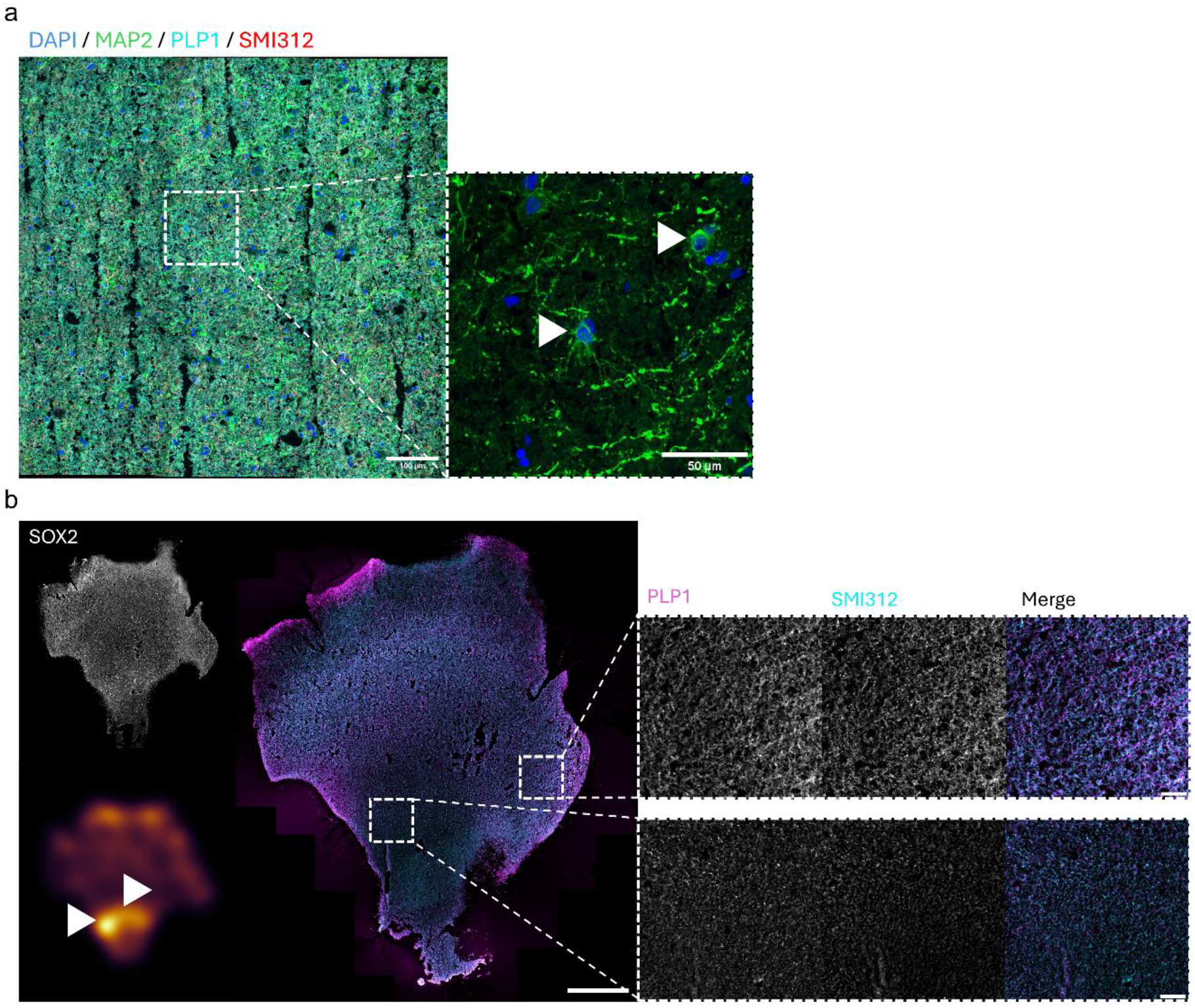
Visualization of MAP2+ neurons and myelinated axons. Fixed-frozen cryosections (12 µm) from a surgical cortex sample of an adult human glioma patient; IDH wild-type tumor. A) Maximum intensity projection 40x confocal image showing a staining for MAP2, PLP1, and SMI312 with a DAPI nuclear counterstain. Zoomed image with arrowheads pointing at MAP2+ cells, indicating their neuronal identity. Scale bars: left 100 µm, right 50 µm. B) Maximum intensity projection 20x image showing a SOX2 stain from figure 2a (upper left) and a Gaussian-weighted inferno colored density map (lower left). The density map is based on the proportion of SOX2+ cells over the total of DAPI+ detected cells, highlighting SOX2+ hotspots. In the density map, black indicates 0 and bright yellow 1 (i.e. hotspot). PLP1 and SMI312 stain (middle) on the same sample with zoomed images at SOX2+ hotspots based on the density map. Zoomed image from a region with a low number of SOX2+ cells (upper) and zoomed image from a region with a high number of SOX2+ cells (lower). Scale bars: 1 mm and 100 µm.

Comparing acute and cultured samples demonstrated a culture-dependent effect on myelin composition and axonal integrity (Fig 2b). This organization of myelin might result from tumor infiltration, axotomy during post-surgical sectioning, or culture-dependent effects on myelinating oligodendrocytes. Consequently, further examination is necessary.

### Oligodendrocyte lineage visualization

Next, to explore the integration of glioma cells into the structural white matter network a staining for cells within the oligodendrocyte lineage was performed (Fig 4). The combination of the SOX10, BCAS1, and ASPA antibodies was used to label the oligodendrocyte lineage, pre-myelinating oligodendrocytes, and mature oligodendrocytes, respectively [27,28]. Focused examination of certain SOX10+ASPA+ cells revealed fragmentation, plausibly linked to biological causes (Fig 4a). Additionally, patient samples included cells positive for both SOX10 and BCAS1, exhibiting characteristic ramified morphology indicative of ongoing myelination processes [27] (Fig 4b). SOX10+BCAS1+ cells were primarily located in grey matter regions and might be independent of glioma infiltration [27].

**Figure 4.**
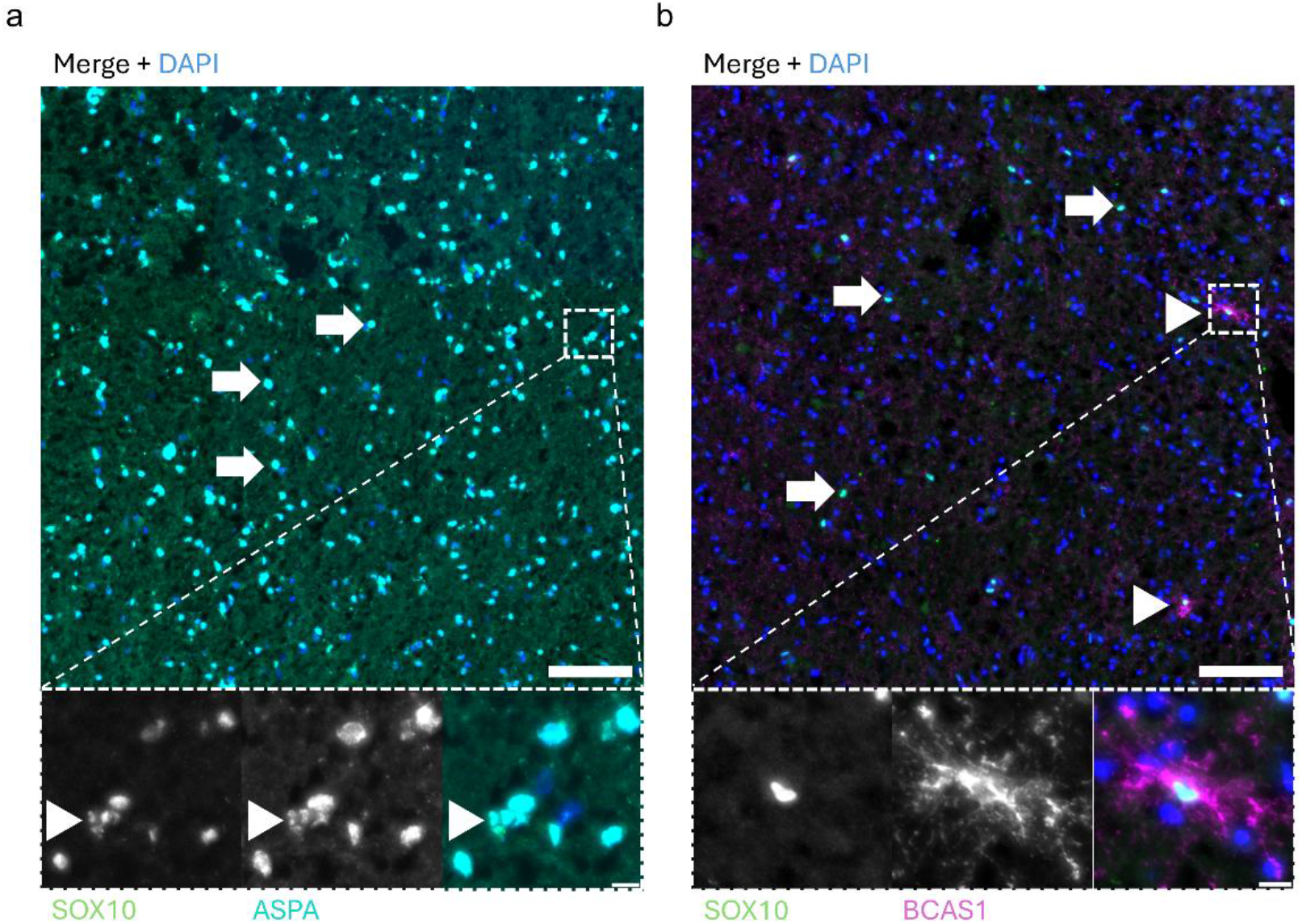
Oligodendrocyte lineage visualization. Maximum intensity projection images from a 20x Slide scanner with fixed-frozen cryosections from a surgical cortex sample of an adult human glioma patient. A) IDH wild-type sample (12 µm). Showing a SOX10 and ASPA stain with a DAPI nuclear counterstain. The image shows examples of clear SOX10+ASPA+ cells (arrows) and examples of fragmented cells (arrowheads). Scale bars: 100 µm and 10 µm. B) IDH mutant sample (20 µm). Shows a SOX10 and BCAS1 stain with a DAPI nuclear counterstain. The image shows SOX10+ cells (arrow) and SOX10+BCAS1+ cells (arrowheads). Scale bars: 100 µm and 10 µm.

Collectively, the observations from immunohistochemistry analyses confirm our ability to visualize glioma infiltration, the neuronal network, myelin coverage, and cells from the oligodendrocyte lineage. However, inter- and intra-sample heterogeneity as well as cultivation-dependent effects were evident, underlining the need for further data collection and comparisons over different culture durations.

## Discussion

Here, an experimental framework was established to analyze glioma cellular networks and their relation to Karnofsky Performance Status (KPS) in a human organotypic slice model. Five human organotypic cortex samples were successfully cultured and stained. Using this framework, glioma infiltration, neurons, myelin, and cells from the oligodendrocyte lineage were successfully visualized after 7 days of culturing. Cultivation-dependent effects were found for neuron count and myelin organization. However, preliminary analyses indicate the feasibility of combining immunohistochemical observations from organotypic slices with patients’ functional status in terms of KPS. Altogether, these observations imply that human organotypic slices are a suitable model for the investigation of glioma functional and structural embedding into the brain network, although further optimization may be desirable.

Other studies have found both similar and dissimilar results in terms of cultivation effects, which are likely attributable to culture media composition. For instance, the study by Ravi [20] found a cultivation-dependent decrease in NeuN+ cells in their human organotypic cortex model. In contrast, other studies have demonstrated reliable maintenance of neuronal architecture and cortical layers after 2 days of culturing using a similar protocol and human model [24]. Although neuron numbers vary between cortical regions [29], and NeuN does not stain all neuronal subtypes [30], the observed quantity in this study was low. Based on research using human organotypic cortex samples, serum-containing culture media does not seem to be optimal for cell viability. Several studies specifically report using serum-free media to prevent astrocyte-dependent inflammation [20] or to enhance survival of their human organotypic slice cultures [31]. The study by Schwarz [32] showed an improvement of survival and network function in these cultures by using human cerebrospinal fluid (hCSF). However, culturing for less than 7 days might not require hCSF [32]. Notably, in 70% of studies employing an organotypic slice model for glioblastoma (GBM) research, serum-free media were used [19]. Together, our study and work of others indicate that human organotypic slices from glioma patients can be used as a reliable model for the study of glioma cellular features, although care should be taken to choose the right medium composition and cultivation time.

Besides optimization of the culturing medium composition and culture duration, effects of resection on the brain tissue should additionally be reconsidered. Any acute damage to the brain, such as post-surgical collection and sectioning of samples, can induce a wound-healing response including reactive gliosis [33,34]. For example, in a damaged region, a loss of oligodendrocytes is observed within the first week. Since reactive gliosis involves the proliferation of astrocytes [33], comparing astrocyte numbers and proliferation in acute and cultured tissue could offer further insight into the post-surgical mechanical stress response. Additionally, the impact of reactive gliosis on both oligodendrocyte cell viability and myelin integrity should be taken into consideration. Furthermore, the immunoreactivity for the SMI312 antibody, directed against the phosphorylated medium and heavy chain, and the absence of this immunoreactivity for the NF-200 antibody, targeting the heavy chain, indicate the specificity for staining a neurofilament state. Phosphorylation of neurofilament was initially considered an indicator of stable, mature axons [26], but later research suggests it may also signify axonal injury [35]. The patchy staining pattern of SMI312 and the unsuccessful NF-200 stain could imply an intermediate or heterogeneous state of intact and compromised axons. This intermediate state of compromised axons is likely a consequence of axotomy induced by tissue processing. However, compromised axons resulting from glioma cell infiltration cannot be excluded. Collectively, post-surgical processing of the samples induces damage-restoring responses, which entail that the duration of culturing determines the time point within the wound-healing response timeline and should be further assessed.

Overall, our successful staining of glioma cells as well as several resident brain cell populations in organotypic cortex from glioma patients will allow future investigations to focus on the organization of structural and functional tumor-brain networks and their relationship to functional status.

## Author contributions

L.R.W. conducted human organotypic cortex slice culturing, followed by processing of cultured slices for immunohistochemistry and immunohistochemistry experiments. L.B. performed image acquisition and part of the immunohistochemistry. L.R.W. and L.B. designed the experiments and analysis. L.R.W. analysed and prepared images for quantification and visualization. M.V.L. arranged patient inclusion and tissue collection. P.W.H., D.N., N.V., and M.B. performed neurosurgeries and provided tissue samples. L.R.W. and D.A.M. wrote the manuscript. L.D. and L.B. and D.A.M. supervised the project.

## Code availability

Code used for data analysis, adapted from QuPath, is publicly available in the <u>project repository</u>.

## Acknowledgements

Special thanks to the department of Integrative Neurophysiology from the Centre for Neurogenomics and Cognitive Research for their help with tissue collection and processing.

## References

1. IJzerman-Korevaar M, Snijders TJ, De Graeff A, Teunissen SCCM, De Vos FYF. Prevalence of symptoms in glioma patients throughout the disease trajectory: a systematic review. J Neurooncol. 2018;140(3):485–496.

2. Douw L, Reijneveld JC, Mandal AS. Multiscale network perspectives on glioma: From tumor biology to symptoms, survival and treatment. Nat Rev Neurol. 2025;22(2):73–89.

3. Zimmermann ML, Van Lingen MR, Koderman E, Dam SC, Breedt LC, Maas DA, et al. Structural network embedding governs peritumor and distant pathological brain activity in glioblastoma. medRxiv [Preprint]. 2026 [posted 2026 May 5; cited 2026 May 21]. Available from: https://doi.org/10.64898/2026.05.05.26352433 doi: 10.64898/2026.05.05.26352433

4. Vargas LO, Latzman S, Tickoo L, Abikenari M, Syed SA, Mittelman L, et al. Gliomas as network diseases: Neuron-tumor interactions, connectome disruption, and clinical implications. Clin Neurol Neurosurg. 2026;267:109475.

5. Osswald M, Jung E, Sahm F, Solecki G, Venkataramani V, Blaes J, et al. Brain tumor cells interconnect to a functional and resistant network. Nature. 2015;528(7580):93–98.

6. Hausmann D, Hoffmann DC, Venkataramani V, et al. Autonomous rhythmic activity in glioma networks drives brain tumor growth. Nature. 2023;613:179–186.

7. Venkataramani V, Tanev DI, Strahle C, Studier-Fischer A, Fankhauser L, Kessler T, et al. Glutamatergic synaptic input to glioma cells drives brain tumor progression. Nature. 2019;573(7775):532–538.

8. Barron T, Yalçın B, Su M, Byun YG, Gavish A, Shamardani K, et al. GABAergic neuron-to-glioma synapses in diffuse midline gliomas. Nature. 2025;639(8056):1060–1068.

9. Tetzlaff SK, Reyhan E, Layer N, Bengtson CP, Heuer A, Schroers J, et al. Characterizing and targeting glioblastoma neuron-tumor networks with retrograde tracing. Cell. 2024;188(2):390-411.e36.

10. Drexler R, Yalçın B, Mancusi R, Rogers A, Shamardani K, Woo PJ, et al. Serotonergic neuron-glioma interactions drive high-grade glioma pathophysiology. bioRxiv [Preprint]. 2025 [posted 2025 Dec 10; cited 2026 May 21]. Available from: https://doi.org/10.64898/2025.12.10.693579 doi: 10.64898/2025.12.10.693579

11. Venkatesh HS, Morishita W, Geraghty AC, Silverbush D, Gillespie SM, Arzt M, et al. Electrical and synaptic integration of glioma into neural circuits. Nature. 2019;573(7775):539–545.

12. Taylor KR, Monje M. Neuron-oligodendroglial interactions in health and malignant disease. Nat Rev Neurosci. 2023;24(12):733–746.

13. Cuddapah VA, Robel S, Watkins S, Sontheimer H. A neurocentric perspective on glioma invasion. Nat Rev Neurosci. 2014;15(7):455–465.

14. Nebeling FC, Fuhrmann F, Mittag M, Musacchio F, Antony H, Gockel N, et al. Microglia-glioblastoma crosstalk mediates glioblastoma invasion at the far infiltration zone. Immunity. 2026;59(4):1075-1091.e4.

15. Mikolajewicz N, Zhai K, Puri A, Miletic P, Tatari N, Wei J, et al. Reactive oligodendrocytes promote glioblastoma progression through CCL5/CCR5-mediated glioma stem cell maintenance. Neuron. 2026;114(2):237-249.e10.

16. Brooks LJ, Clements MP, Burden JJ, Kocher D, Richards L, Devesa SC, et al. The white matter is a pro-differentiative niche for glioblastoma. Nat Commun. 2021;12:2184.

17. Hu J, Bao H, Liu X, Fang S, Yan Z, Wang Z, et al. Glioma-white matter tract interactions: A diffusion magnetic resonance imaging-based 3-tier classification and its clinical relevance. Neuro Oncol. 2025;27(7):1888–1898.

18. Huszthy PC, Daphu I, Niclou SP, Stieber D, Nigro JM, Sakariassen PO, et al. In vivo models of primary brain tumors: pitfalls and perspectives. Neuro Oncol. 2012;14(8):979–993.

19. Petralia CCT, D’Amico AG, D’Agata V, Broggi G, Barbagallo GMV. Ex vivo organotypic brain slice models for glioblastoma: a systematic review. Cancers (Basel). 2026;18(3):372.

20. Ravi VM, Joseph K, Wurm J, Behringer S, Garrelfs N, Errico PD, et al. Human organotypic brain slice culture: a novel framework for environmental research in neuro-oncology. Life Sci Alliance. 2019;2(4):e201900305.

21. Louis DN, Perry A, Wesseling P, Brat DJ, Cree IA, Figarella-Branger D, et al. The 2021 WHO Classification of Tumors of the Central Nervous System: a summary. Neuro Oncol. 2021;23(8):1231–1251.

22. Chaichana K, Parker S, Olivi A, Quiñones-Hinojosa A. A proposed classification system that projects outcomes based on preoperative variables for adult patients with glioblastoma multiforme. J Neurosurg. 2010;112(5):997–1004.

23. Wilbers R, Metodieva VD, Duverdin S, Heyer DB, Galakhova AA, Mertens EJ, et al. Human voltage-gated Na+ and K+ channel properties underlie sustained fast AP signaling. Sci Adv. 2023;9(41):eade3300.

24. Ting JT, Kalmbach B, Chong P, De Frates R, Keene CD, Gwinn RP, et al. A robust ex vivo experimental platform for molecular-genetic dissection of adult human neocortical cell types and circuits. Sci Rep. 2018;8(1):8407.

25. Trojanowski J, Walkenstein N, Lee V. Expression of neurofilament subunits in neurons of the central and peripheral nervous system: an immunohistochemical study with monoclonal antibodies. J Neurosci. 1986;6(3):650–660.

26. Ulfig N, Nickel J, Bohl J. Monoclonal antibodies SMI 311 and SMI 312 as tools to investigate the maturation of nerve cells and axonal patterns in human fetal brain. Cell Tissue Res. 1998;291(3):433–443.

27. Fard MK, Van Der Meer F, Sánchez P, Cantuti-Castelvetri L, Mandad S, Jäkel S, et al. BCAS1 expression defines a population of early myelinating oligodendrocytes in multiple sclerosis lesions. Sci Transl Med. 2017;9(419):eaam7816.

28. Mironova YA, Dang B, Heo D, Xu YKT, Hsu AY, Von Bernhardi JE, et al. Myelin is repaired by constitutive differentiation of oligodendrocyte progenitors. Science. 2026;391(6783):eadu2896.

29. Von Bartheld CS, Bahney J, Herculano-Houzel S. The search for true numbers of neurons and glial cells in the human brain: a review of 150 years of cell counting. J Comp Neurol. 2016;524(18):3865–3895.

30. Lyck L, Santamaria ID, Pakkenberg B, Chemnitz J, Schrøder HD, Finsen B, et al. An empirical analysis of the precision of estimating the numbers of neurons and glia in human neocortex using a fractionator-design with sub-sampling. J Neurosci Methods. 2009;182(2):143–156.

31. Eugène E, Cluzeaud F, Cifuentes-Diaz C, Fricker D, Duigou CL, Clemenceau S, et al. An organotypic brain slice preparation from adult patients with temporal lobe epilepsy. J Neurosci Methods. 2014;235:234–244.

32. Schwarz N, Hedrich UBS, Schwarz H, Harshad PA, Dammeier N, Auffenberg E, et al. Human cerebrospinal fluid promotes long-term neuronal viability and network function in human neocortical organotypic brain slice cultures. Sci Rep. 2017;7(1):12249.

33. Burda JE, Sofroniew MV. Reactive gliosis and the multicellular response to CNS damage and disease. Neuron. 2014;81(2):229–248.

34. Bak A, Koch H, Van Loo KM, Schmied K, Gittel B, Weber Y, et al. Human organotypic brain slice cultures: a detailed and improved protocol for preparation and long-term maintenance. J Neurosci Methods. 2024;404:110055.

35. Petzold A. Neurofilament phosphoforms: surrogate markers for axonal injury, degeneration and loss. J Neurol Sci. 2005;233(1-2):183–198.

